# Differential intra-host infection kinetics in *Aedes aegypti* underlie superior transmissibility of African relative to Asian Zika virus

**DOI:** 10.1101/2023.05.24.542190

**Authors:** Rinyaporn Phengchat, Phonchanan Pakparnich, Chatpong Pethrak, Jutharat Pengon, Channarong Sartsanga, Nunya Chotiwan, Kwanchanok Uppakara, Kittitat Suksirisawat, Louis Lambrechts, Natapong Jupatanakul

## Abstract

Despite numerous studies highlighting the higher transmissibility of the African Zika virus (ZIKV) lineage compared to the Asian lineage in mosquito vectors, little is known about how the viruses interact with different tissues during the early steps of mosquito infection. To address this gap, we aimed to characterize intra-host infection barriers by combining a tissue-level monitoring of infection using plaque assays and a novel quantitative analysis of single-cell level infection kinetics by *in situ* immunofluorescent staining. Our results revealed that, in *Aedes aegypti*, an African ZIKV strain exhibited a higher replication rate across various tissues than an Asian ZIKV strain. This difference was potentially due to a higher virus production in individual cells, faster spread within tissues, or a combination of both factors. Furthermore, we observed that higher bloodmeal titers resulted in a faster viral spread to neighboring cells suggesting that intra-host infection dynamics depend on inoculum size. We also identified a significant bottleneck during midgut infection establishment for both ZIKV lineages, with only a small percentage of the virus population successfully initiating infection. Finally, the *in situ* immunofluorescent staining technique enabled the examination of virus infection characteristics in different cell types and revealed heterogeneity in viral replication. Together, these findings demonstrate that differences in intra-host infection kinetics underlie differential transmissibility between African and Asian ZIKV lineages. This information could serve as a starting point to further investigate the underlying mechanisms and ultimately inform the development of alternative control strategies.

**Importance:** The recent Zika virus (ZIKV) epidemic in the Americas highlights its potential public health threat. While the Asian ZIKV lineage has been identified as the main cause of the epidemic, the African lineage, which has been primarily confined to Africa, has shown evidence of higher transmissibility in *Aedes* mosquitoes. To gain a deeper understanding of this differential transmissibility, our study employed a combination of tissue-level infection kinetics and single-cell level infection kinetics using *in situ* immunofluorescent staining. We discovered that the African ZIKV lineage propagates more rapidly and spreads more efficiently within mosquito cells and tissues than its Asian counterpart. This information lays the groundwork for future exploration of the viral and host determinants driving these variations in propagation efficiency.

## Introduction

Zika virus (ZIKV) is a mosquito-borne flavivirus primarily transmitted between humans through bites of infected *Aedes* mosquitoes. First identified in Uganda in 1947 from a sentinel monkey, sporadic ZIKV cases have been reported in Africa and Asia over the following decades (Dick et al., 1952; Liu et al., 2019). ZIKV spread to several Pacific islands between 2013-2014 and reached South America in 2015, causing an outbreak that affected hundreds of thousands of people, underlining the potential of emerging and re-emerging arthropod-borne (arbo) flavivirus outbreaks (Liu et al., 2019). Phylogenetic analysis of ZIKV genome sequences revealed two major ZIKV lineages, the African and the Asian lineage, with studies indicating that the African lineage is more transmissible by mosquito vectors than the Asian lineage (Aubry et al., 2021; Bialosuknia et al., 2017; Roundy et al., 2017). However, the complex infection kinetics of ZIKV in mosquitoes and its interactions with individual tissues prior to transmission remain poorly understood.

Following ingestion of an infectious bloodmeal by mosquito vectors, the viruses must propagate in different body compartments and eventually be secreted in mosquito saliva when taking subsequent bloodmeals for transmission to occur. Several anatomical bottlenecks must be overcome for successful virus transmission in insects, including midgut infection, midgut escape, dissemination, salivary gland infection, and salivary gland escape barriers (Franz, 2015). Traditional techniques for studying virus infection kinetics, such as quantitative RT-PCR or infectious virus titration with cell-based assays, involve tissue grinding or lysis to extract and quantify viral RNA or infectious particles. While these methods can provide information on infection level, they have a significant limitation: the loss of spatial information of the infection kinetics. In contrast, *in situ* immunofluorescent staining preserves the spatial information of virus infection. Although this technique has the potential to provide a more comprehensive overview of infection kinetics including the sites of initial infection, cell-type tropism, rate of virus propagation and spread within the tissues, its application has remained limited to investigate arbovirus-mosquito infection.

In this study, we infected *Aedes aegypti* (*Ae. aegypti*) mosquitoes with African (DAK AR41524) or Asian (SV0010/15) ZIKV and investigated intra-host infection kinetics at the tissue level by virus titration, and at the single-cell level using immunofluorescent staining. The results provide evidence that the African strain ZIKV has superior infectivity from the establishment of midgut infection, the spread of infection in the tissue to the replication at the cellular level. Our data demonstrate that the detailed analyses of *in situ* immunofluorescent staining offers a powerful tool to compare steps that determine differential infectivity of arboviruses in the insect vector.

## Materials and Methods

### Ethics statement

This study was carried out in strict accordance with the recommendations in the Guide for the Care and Use of Laboratory Animals of the National Institutes of Health. Mice were used only for mosquito rearing as a blood source, according to the protocol (BT-Animal 05/2564). Mosquito infection assays were performed followed the approved protocol (BT-Animal 05/2564). Both animal protocols were approved by the BIOTEC Committee for use and care of laboratory animals. Human erythrocytes were used for infection by artificial membrane feeding. Blood was collected from human volunteer following the approved protocol (NIRB-052-2563).

### Animal maintenance

A laboratory strain of *Ae. aegypti* obtained from the Department of Medical Sciences, National Institute of Health Thailand and further maintained at BIOTEC’s insectary for an additional 36-38 generations was used for all the infection studies. Mosquitoes were maintained at 28 ± 1 °C with 70% relative humidity and a 12-hour day/night, 30-minute dusk/dawn lighting cycle. The larvae were fed on powdered fish food (Tetra Bits). Adults were fed on a sterile 10% sucrose solution. To obtain the eggs for colony maintenance, mosquitoes were allowed to feed on ICR mice anesthetized with 2% Avertin (2,2,2-Tribromoethanol, Sigma, T48402).

### Virus propagation and titration

The DAK AR41524 strain was used as a representative of the African ZIKV lineage while the SV0010/15 strain was used as a representative of the Asian ZIKV lineage. The *Aedes albopictus* cell line C6/36 (ATCC CRL-1660) was used for amplification of all virus stocks and for *in vitro* infection kinetics. C6/36 cells were maintained at 28 °C in Leibovitz’s L-15 (L-15) medium (Sigma, L4386) with 10% fetal bovine serum (FBS, PAN BIOTECH, P30-3031), 2% tryptose phosphate broth (Sigma, T9157), 1X non-essential amino acids (Gibco, 11140-050), 1X Pen/Strep (100 U/ml of penicillin and 100 μg/ml of streptomycin, Cytiva, SV30010).

Virus stocks were prepared in C6/36 cells according to a previously published protocol (Aubry et al., 2021). After the cells were cultured to 80% confluency in 75 cm^2^ flasks, all supernatants were removed and replaced with the virus stocks at the multiplicity of infection (MOI) of 0.1 in 5 mL incomplete L-15 medium for 2 hours. After virus incubation, supernatant was removed, and replaced with 2% FBS L-15 medium then further incubated at 28 °C. The supernatant was collected at 6-7 days post inoculation then supplemented with FBS to final concentration of 20% and stored at 80 °C until further use.

Plaque assay was used to determine virus titers following a previously published protocol (Jupatanakul et al., 2017). Briefly, 100 μL of virus suspension was added to BHK-21 cells seeded in 24-well plate at 80% confluency. Inoculated plates were then gently rocked at room temperature for 15 minutes before an incubation at 37 °C, 5% CO_2_ for 45 minutes. After incubation, 1 mL of overlay medium (1% methylcellulose (Sigma, M0512) in Minimal Essential Medium (MEM) supplemented with 2% FBS and 1X Pen/Strep) was added to each well then further incubated at 37 °C, 5% CO_2_ for 5 days. The plates were then fixed and stained with 0.5% crystal violet (Sigma, C6158) in 1:1 methanol/acetone fixative for 1 hour at room temperature. Stained plates were then washed under running tap water and air dry before plaque counting.

### Mosquito infection by artificial membrane feeding

Mosquitoes were orally challenged with either the African or the Asian ZIKV strain using the Hemotek artificial membrane feeding system according to a previously published protocol (Aubry et al., 2021). Briefly, 7-day-old female *Ae. aegypti* mosquitoes were deprived of sucrose solution overnight before being offered an artificial infectious blood meal containing 40% washed human erythrocytes and virus stock diluted to desired feeding titers for 30 minutes. After feeding, mosquitoes were anesthetized in a refrigerator for 15 minutes, and fully engorged females were sorted on ice. Blood-fed mosquitoes were maintained in waxed paper cups with 10% sucrose solution in a climate-controlled chamber under controlled insectary conditions as mentioned above.

### Mosquito dissection and salivation assay

Mosquitoes were cold anesthetized in a refrigerator or on ice for 15 minutes before surface sterilization in 70% ethanol for 1 minute followed by two PBS washes. Mosquitoes were then individually dissected in drops of 1X PBS. Midgut, carcasses, and salivary glands were collected in 150 μL of MEM supplemented with 10% FBS and 1X Pen/Strep and stored at -80 °C for further analysis. Tissues were homogenized using 0.5 mm glass beads with Bullet Blender Tissue Homogenizer (NextAdvance). The 10-fold serially diluted samples were titrated by plaque assay as described above. Mosquito saliva was collected according to a previously published protocol (Aubry et al., 2021). Briefly, mosquitoes were paralyzed with triethylamine before inserting the proboscis into a pipette tip containing 20 μL of MEM supplemented with 10 % FBS and 1X Pen/Strep. After 45 minutes of salivation, the medium in the tips were mixed with 180 μL of MEM supplemented with 2% FBS and 1X Pen/Strep and immediately titrated by plaque assay mosquito. Mosquito salivary glands, midgut and carcasses were dissected from anesthetized mosquitoes and stored at -80 °C for virus titration by plaque assay.

### Immunofluorescent staining of infected midguts

Dissected midguts were cut in half to remove blood bolus then fixed with 4% paraformaldehyde in PBS for 2 hours at room temperature and permeabilized using 2% Triton X-100 in PBS for 15 minutes. After permeabilization, midguts were incubated in 2% bovine serum albumin in PBS for 1 hour. Midguts were incubated with primary antibody anti-flavivirus envelope 4G2 antibody produced in-house in 1X PBS at 4 °C overnight followed by 1:500 secondary antibody anti-mouse IgG Alexa 488 (Invitrogen, A28175) in 1X PBS and Hoechst33342 (Invitrogen, 62249) for 3 hours at room temperature. Midguts were mounted onto glass slides in Vectashield Plus (Vector Laboratories, H1900). Images were taken at the same fluorescent settings for both ZIKV strains under an inverted fluorescence microscope (Olympus IX81) and laser scanning confocal microscopes (Nikon AXR and Carl Zeiss LSM900) using 20X objective lens (CFI Apochromat LWD Lambda S 20XC WI, LD Plan-Neofluar 20X/0.4 Corr M27). The z-section distance was 2.5 μm and the total z-section thickness was set between 30–60 μm. Z-section projection with maximum intensity was done using Fiji version 1.53t (Schindelin et al., 2012). The whole midgut tissues were used for cell-to-cell infection kinetics analyses.

### Infected cell count and fluorescent intensity analysis

Fiji and CellProfiler 4.2.1 (Stirling et al., 2021, http://www.cellprofiler.org; Broad Institute, Cambridge, MA) were used for image analysis. In Fiji, grayscale images of infected cells and nuclei were proceeded to background subtraction and object segmentation to generate binary images using following plugins: Trainable Weka Segmentation (TWS) (Arganda-Carreras et al., 2017) for infected cells segmentation and StarDist 2D (Schmidt et al., 2018) for nuclei segmentation. Watershed and segmented line tools in Fiji were applied to binary images of infected cells generated from TWS to separate individual cells.

Workflows in CellProfiler were created to count the number of infected foci, identify the number of cells in infected foci and measure fluorescent intensity of infected cells and their nuclei. To count the number of infected cells in foci, binary images of infected foci and nuclei were used as input images for identifying primary objects. While to measure fluorescent intensity, binary images of individual infected cells and nuclei were used as input images. A mask of infected foci/cells was used for extracting nuclei of infected cells. Additional details on the image analysis can be found in the Supplementary Figure S1. Fluorescent intensity of 4G2 staining was measured from maximum Z-projected images of infected midguts taken with confocal microscopes with the 20X objective lens. Only infected cells from midgut monolayer and non-overlapping infected cells from midgut multilayer were measured.

### Data availability

The data and immunofluorescence images from this study are available upon request to the corresponding author.

## Results

### The African ZIKV strain displays faster intra-host infection kinetics than the Asian ZIKV strain

To identify intra-host infection barriers that lead to different transmissibility between African and Asian ZIKV lineages, we compared infection kinetics of two representative strains during midgut infection, dissemination (virus escape from the midgut and systemic infection), salivary gland infection, and release in saliva over time. A laboratory strain of Thai *Ae. aegypti* was fed with approximately 7 log_10_ PFU/mL of African (DAK AR41524) or Asian ZIKV (SV0010/15). Uncontrolled variation in infectious dose was less than 0.2 log_10_ PFU/mL across experiments and ZIKV strains.

The infection kinetics revealed that African ZIKV was more efficient at establishing midgut infection than Asian ZIKV (Figure 1). Over 50% of mosquitoes fed with African ZIKV had detectable infectious virus in their midguts at 1 day post infectious blood meal (dpibm), while no detectable infection was observed for Asian ZIKV. Although midgut viral titers of the African strain remained higher than those of the Asian strain at 2-3 dpibm, midgut infection level eventually plateaued, and the Asian strain reached comparable titers to the African strain from 4 dpibm onward.

**Figure 1.**
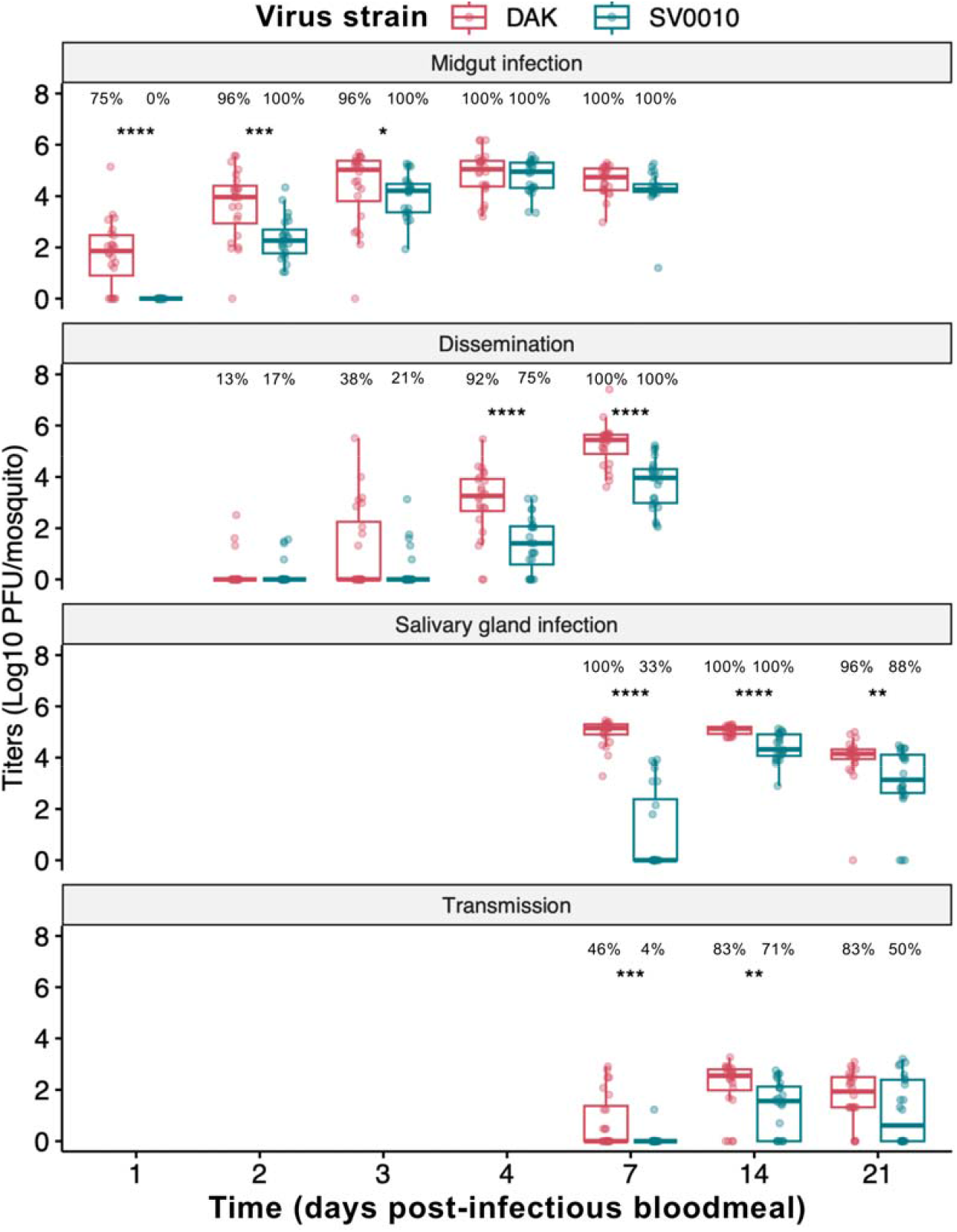
African ZIKV has more rapid intra-host infection kinetics than Asian ZIKV. Intra-host infection kinetics of African (DAK) and Asian (SV0010) ZIKV strains in *Ae. aegypti*. The midgut infection was investigated at 1, 2, 3, 4, and 7 days post-infectious bloodmeal (dpibm). Dissemination (escape of virus from the midgut and replication in the body) was measured in the carcasses at 2, 3, 4, and 7 dpibm. Salivary gland infection and transmission (virus titers in saliva) was measured at 7, 14, and 21 dpibm. Data was summarized from two blood feeding experiments with at least 10 mosquitoes from each. The feeding titers for the midgut infection and dissemination samples were 7.2-7.38 log_10_ PFU/mL for DAK and 7.34-7.38 log_10_ PFU/mL for SV0010. Box and scatter plots represent the distribution of viral titers in log_10_-transformed PFU/mosquito. Statistical analysis comparing infection level was conducted using Kruskal-Wallis followed by Dunn’s posthoc test in R. *: p<0.05, **: p<0.01, ***: p<0.001, ****: p<0.0001. Percentage on top of each graph indicate infection prevalence (number of samples with positive virus x 100/number of total sample).

During the subsequent phase of intra-host infection kinetics, similar dissemination prevalence (% of mosquitoes with detectable virus outside the midgut) at early timepoints was observed for both strains, suggesting that midgut escape may not be a bottleneck in the Asian ZIKV transmission cycle (Figure 1). However, African ZIKV displayed significantly higher dissemination titers at 4 dpibm, and this trend persisted through 7 dpibm. The higher virus titers of African ZIKV during later timepoints of dissemination might be due to a higher number of viruses available to escape the midgut to initiate a systemic infection, or a higher replication rate of the African ZIKV strain in secondary organs compared to the Asian ZIKV strain.

Next, we investigated salivary gland infection and transmission kinetics at 7, 14, and 21 dpibm. We found that the African ZIKV strain reached the salivary glands earlier and replicated more efficiently than the Asian ZIKV strain (Figure 1). African ZIKV reached the salivary glands of all mosquitoes by 7 dpibm, while Asian ZIKV was only detected in the salivary glands of 33% of blood-fed mosquitoes. Although salivary glands of all mosquito blood-fed with Asian ZIKV became infected by 14 dpibm, average virus titers of African ZIKV were higher than those of Asian ZIKV across all time points measured suggesting a more robust virus propagation in salivary gland tissue.

African ZIKV could be detected in saliva from 46% of mosquito at 7 dpibm while only one out of 24 of the Asian ZIKV blood-fed mosquitoes had detectable infectious virus in saliva (Figure 1). The prevalence and titers in saliva of both viruses increased from 7 to 14 dpibm. Interestingly, the saliva virus titers at 21 dpibm was lower than 14 dpi for both viruses suggesting lower transmission efficiency at the very late timepoint. These results confirm the higher transmissibility of the African ZIKV strain than the Asian ZIKV strain in *Ae. aegypti* in our study system. Taken together, our results suggested that the higher transmission efficiency of the African ZIKV strain was a result from a more rapid infection starting from the midgut infection, dissemination, and salivary gland infection. The lower transmissibility of the Asian ZIKV strain may therefore be caused by the lower amount of virus that is available to escape salivary glands given the lower prevalence and titers of salivary gland infection.

Infection kinetics in all mosquito tissues revealed that the key feature of the African ZIKV strain was a higher replication rate in infected cells in various tissue types prior to salivary gland escape. However, it remains unclear whether the increased replication is due to higher virus production in individual cells, faster viral spread within tissues, or both. The results shown in Figure 1 suggest that the replication kinetics of two ZIKV strains can be readily differentiated during the establishment of midgut infection. Our experimental approach allows the initial viral inoculum to be controlled (by using the same titer in the bloodmeal) so that differential cell-to-cell spread can easily be observed in the intact tissue. We next investigated the cell-to-cell level infection kinetics in the midgut by focusing on the three critical steps for successful midgut infection: i) the establishment of virus infection in midgut epithelial cells (primary infection), ii) the replication of virus in the primary infected cells, and iii) the spread of virus from the primary infected cells to neighboring cells (secondary infection).

### African ZIKV exhibits superior primary midgut infection success compared to Asian ZIKV

To compare the infectivity of African and Asian ZIKV during primary infection of the mosquito midgut, we compared the number of midgut cells that became infected after ingestion of an infectious blood meal containing approximately 7 log_10_ PFU/mL of each ZIKV strain. Infected cells in the midgut at 1 dpibm were visualized using immunofluorescent staining. The number of infected cell foci for the African ZIKV strain was more than twice that of the Asian ZIKV strain, with a mean number of 279±322 and 125±53 infection foci per midgut, respectively (Figure 2). These results indicate that, given similar blood meal titers, African ZIKV establishes midgut infection more effectively than Asian ZIKV.

**Figure 2.**
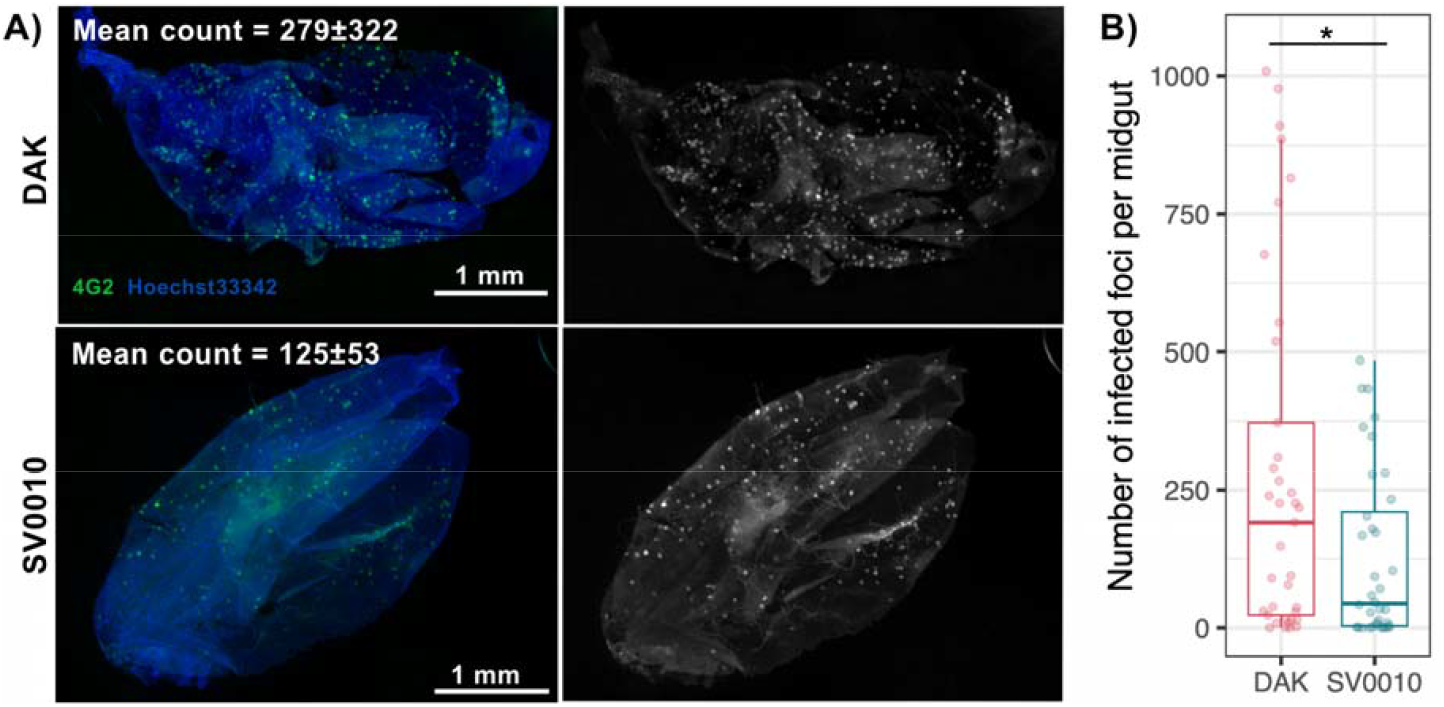
African ZIKV establishes midgut infection better than Asian ZIKV. Left) Representative image of *Ae. aegypti* midguts infected by the Asian ZIKV strain (SV0010) or the African ZIKV strain (DAK) at 1 dpibm. Right) Box and scatter plot comparing number of infected cell foci in each midgut. Each dot represent data from an individual midgut. Data was summarized from two blood feeding experiments with at least 10 midguts from each. The feeding titers for the midgut infection and dissemination samples were 7.2-7.38 log_10_ PFU/mL for DAK and 7.34-7.38 log_10_ PFU/mL for SV0010. Statistical analysis was conducted using student t-test, *: p<0.01.

### Midgut cells infected with African ZIKV display stronger immunofluorescence staining than with Asian ZIKV

To further investigate ZIKV infection dynamics, we assessed the intensity of immunofluorescence staining in each infected cell (Figure 3). In total, we examined 1,417 infected cells for the African ZIKV strain and 318 infected cells for the Asian ZIKV strain. Our results showed that the normalized immunofluorescence intensity in midgut cells infected by the African ZIKV strain was significantly higher than that in cells infected by the Asian ZIKV strain (mean normalized signals of 1.36 ±0.50 and 0.80±0.33, respectively, p<0.0001). The stronger immunofluorescence staining in infected cells suggests that at 1 dpibm, a larger amount of ZIKV E proteins were produced in cells infected with the African strain compared to those infected with the Asian strain. This could be attributed to either a higher level of virus production per cell or a more rapid cellular replication cycle.

**Figure 3:**
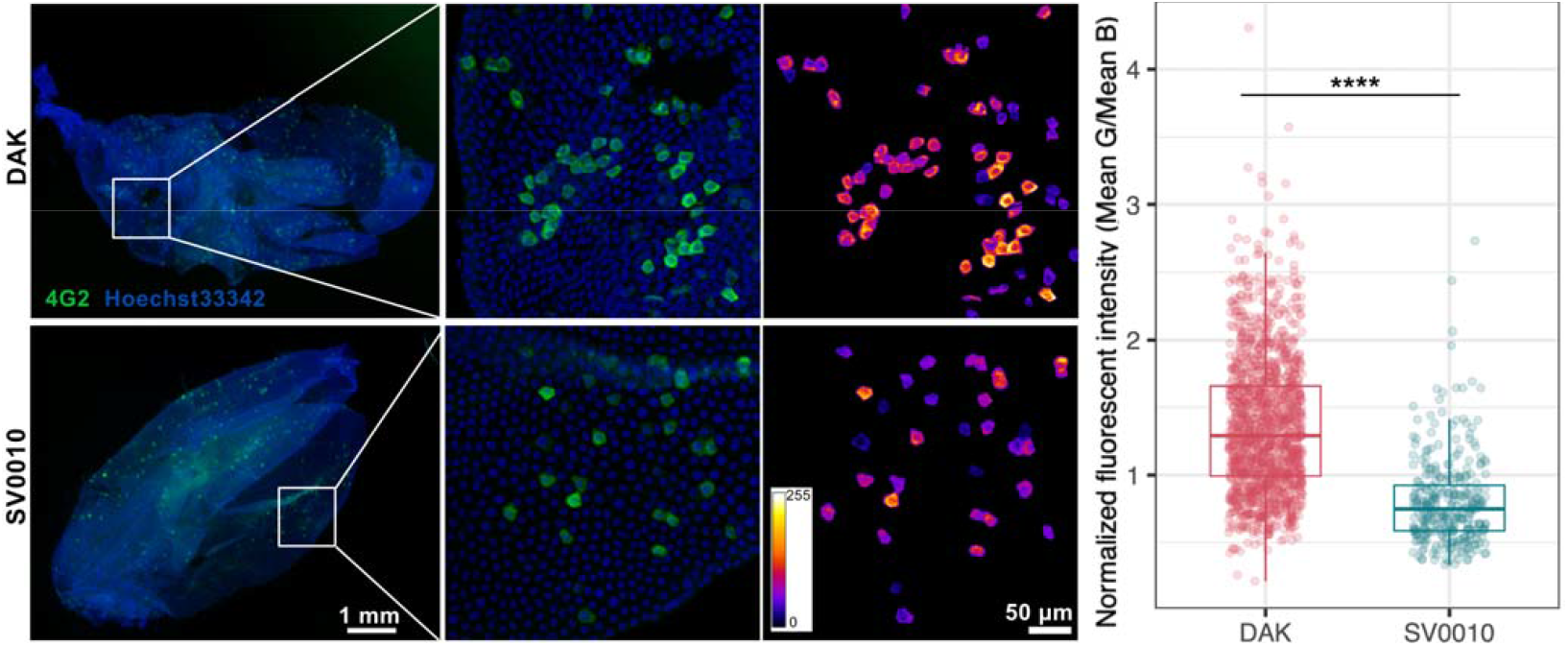
Midgut cells infected with African ZIKV display stronger immunofluorescence signals than with Asian ZIKV. (Left) Representative images of *Ae. aegypti* midguts infected by the Asian ZIKV strain (SV0010) or the African ZIKV strain (DAK) ZIKV at 1 dpibm. The overall midgut images were epifluorescence images under a 4X objective lens. The zoomed images were maximum projected images of confocal z-stacks under a 20X objective lens. The heatmaps demonstrate the green channel fluorescence intensity on a scale ranging from 1 to 255 arbitrary units. (Right) Box and scatter plots comparing immunofluorescence intensity of each infected cell between the two ZIKV strains. Whole mosquito midguts were used for immunofluorescence intensity analyses. The normalized fluorescence intensity was calculated by dividing mean Alexa 488 fluorescence intensity of each infected cell with mean Hoechst 33342 fluorescence intensity of its nucleus. Because the tissue can be folded or have multiple layers in the *in situ* analysis, infected cells in the areas having overlapped infected regions were excluded to avoid inaccurate evaluation of fluorescence intensity of each cell. Each dot represents data from each individual infected cell. Data was summarized from two blood feeding experiments with at least 5 midguts from each. The feeding titers for the midgut infection and dissemination samples were 7.2-7.38 log_10_ PFU/mL for DAK and 7.34-7.38 log_10_ PFU/mL for SV0010. Statistical analysis was conducted using student t-test, ****: p<0.0001.

### African ZIKV spreads to neighboring midgut cells faster than Asian ZIKV

The later stage of tissue-level infection kinetics during midgut infection involves the spread of infection from primary infected cells to neighboring cells (secondary infection). In this experiment, we compared the rate of virus spread in the midgut tissue by measuring the size of infection foci (number of infected cells in each infection focus) at 1 dpibm. With bloodmeal titers of 7 log_10_ PFU/mL, we observed that at 1 dpibm, mosquito midguts infected with African ZIKV displayed a lower percentage of infection foci containing only a single infected cell (primary infection) compared to Asian ZIKV (76.41% and 86.35%, respectively; Figure 4). Notably, the number of foci with more than one infected cell (2, 3, 4, and >4 infected cells per focus) was significantly higher for the African strain than for the Asian strain, indicating faster secondary infection and a greater spread to neighboring cells of the African strain (Figure 4).

**Figure 4:**
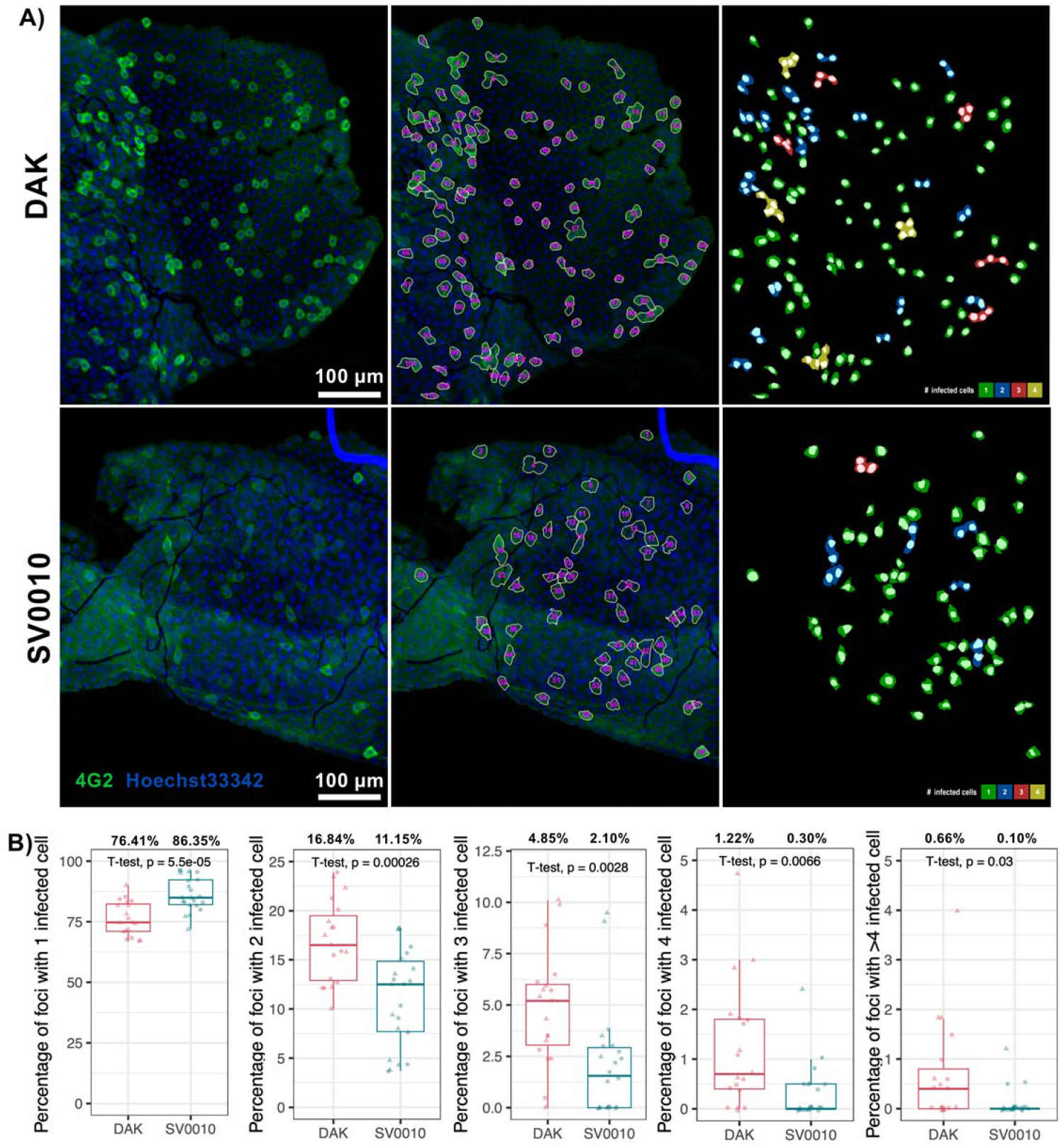
African ZIKV has more foci with secondary infection than Asian ZIKV. A) Representative images demonstrating the infected cell counting. Left: Projection of representative z-stacked images. Middle: segmentation of each focus of infected cells. Some areas of the midgut were folded on top of each other thus having more than one layer of epithelial cells. Right: color-coding representing number of infected cells in each focus. B) Box and scatter plots comparing the percentage of infection foci with 1, 2, 3, 4, or >4 infected cells between midguts infected with Asian (SV0010) or African (DAK) ZIKV strains at 1 dpibm. Data was summarized from two blood feeding experiments with at least 10 midguts from each. The feeding titers for the midgut infection and dissemination samples were 7.2-7.38 log_10_ PFU/mL for DAK and 7.34-7.38 log_10_ PFU/mL for SV0010. Each dot represents data from an individual midgut. Statistical analysis was conducted using student t-test, ****: p<0.0001.

In addition to comparing the size of infection foci at 1 dpibm, we conducted an experiment to examine the expansion of infection foci over time (Figure 5). Given that bloodmeal titers of 7 log_10_ PFU/mL resulted in hundreds of primary infected cells, measuring the expansion of infected cell foci during subsequent timepoints with such a high number of primary infections was infeasible (Supplementary Figure S2). Therefore, bloodmeal titers of approximately 5 log_10_ PFU/mL were used to infect *Ae. aegypti* in this experiment. To measure the expansion of infection foci, mosquito midguts were collected at 1, 2, and 3 dpibm for immunofluorescence staining. We quantified the number of infected cells within infection foci and calculated the percentage of foci containing 1, 2-4, 5-8, 9-20, 21-50, or ≥50 infected cells per focus. Interestingly, with the bloodmeal titers of 5 log_10_ PFU/mL, none of the infected foci had any secondary infection for either African or Asian ZIKV on the first day (Figure 5). This is in contrast with the results obtained using higher bloodmeal titers of 7 log_10_ PFU/mL, which resulted in secondary infection observed at 1 dpibm (Figure 4). Nonetheless, we found that African ZIKV spread to neighboring cells more rapidly than Asian ZIKV. At 2 dpibm, only 13.92% of midgut foci had primary infection (1 infected cell in each focus) for the African ZIKV strain, while midguts still had primary infection in 63.51% of foci for the Asian ZIKV strain. By 3 dpibm, all African ZIKV-infected foci exhibited secondary infection with at least 5 cells per foci, while Asian ZIKV-infected foci still had primary infection in 9.21% of cases and secondary infection in 22.37% of foci with fewer than 4 cells. The faster spread of African ZIKV resulted in larger average infected focus sizes during 2-3 dpibm (Figure 5D). The African ZIKV and Asian ZIKV had average focus sizes of 8±6 and 2±2 infected cells per focus at 2 dpibm, and 10±8 and 29±22 infected cells per focus at 3 dpibm, respectively.

**Figure 5:**
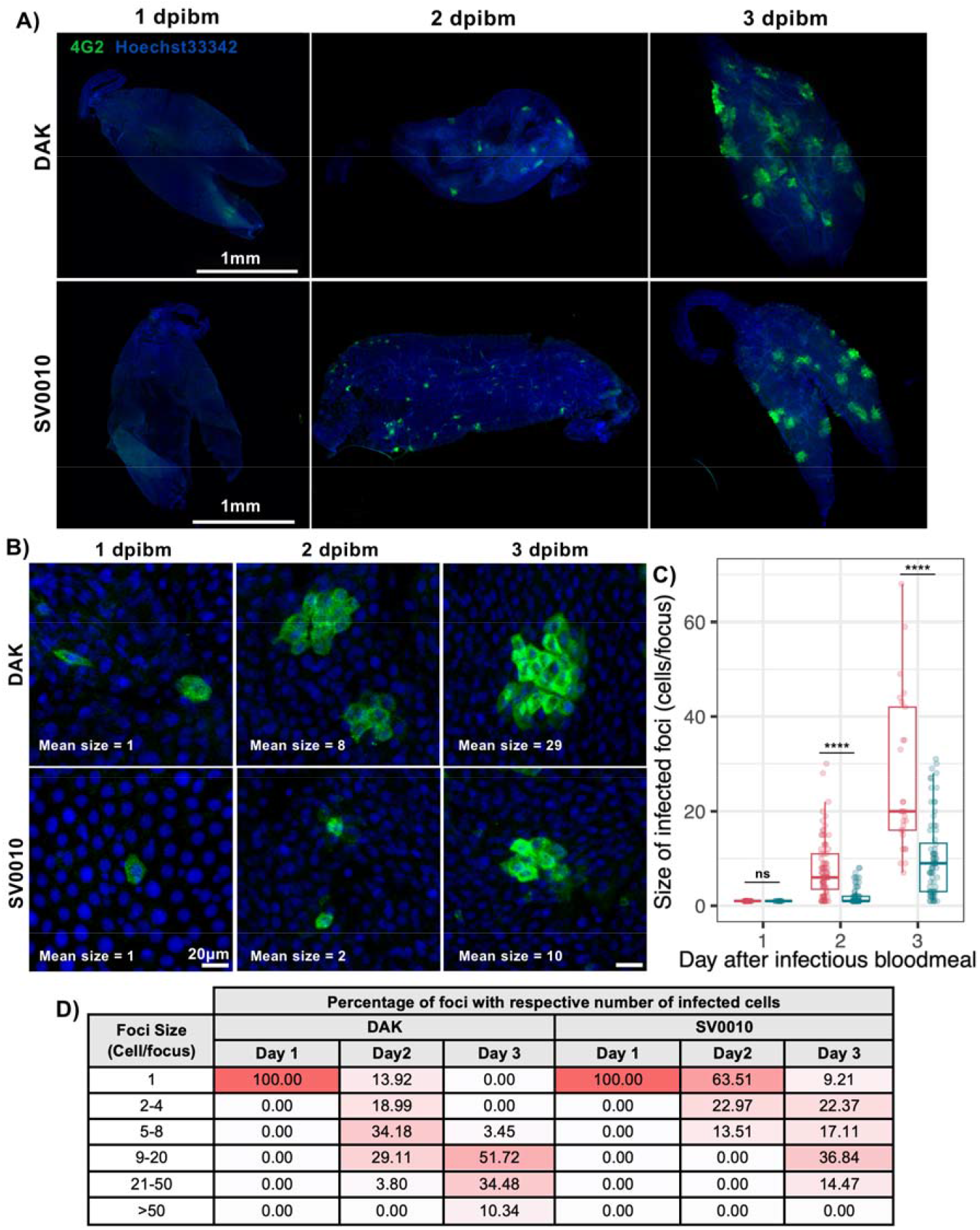
African ZIKV spreads faster to neighboring cells than Asian ZIKV. (A) Overall images of *Ae. aegypti* midguts infected with 5 log_10_ PFU/mL of the African ZIKV strain (DAK) or the Asian ZIKV strain (SV0010) at 1, 2 and 3 dpibm. (B) Higher magnification representative images (200X) demonstrating the size of infection foci. (C) Box and scatter plots demonstrating the number of cells in each infection focus of African and Asian ZIKV. The feeding titers for the midgut infection and dissemination samples were 4.79-4.84 log_10_ PFU/mL for DAK and 5.14-5.25 log_10_ PFU/mL for SV0010. Statistical analysis was conducted using student t-test, ****: p<0.0001. (D) Table comparing the distribution of infection focus sizes over time during 1, 2, and 3 dpibm.

### Progression of ZIKV infection in the midgut tissue varies between different midgut cell populations

The midgut tissue consists of distinct cell populations with varying morphologies and functions. Based on previous research, cells with large cytoplasm and nuclei are likely enterocytes (ECs) while cells with smaller nuclei are likely enteroendocrine cells (EEs), and undifferentiated progenitors including intestinal stem cells (ISCs) and enteroblasts (EBs) (Bonfini et al., 2016; Hixson et al., 2021). We observed that all the primary infected cells are likely ECs as all they have large nucleus and cytoplasm. As the infection progressed, we observed cells with intense immunofluorescence staining at the center of each infection focus. These cells were surrounded by cells of similar size but with subtle decrease in immunofluorescence intensity along the radial distance from the center cells (Figure 6A). This indicates that the infection gradually spread from the primary infected cells to neighboring ECs. Interestingly, in some infected foci, we observed the infection in small satellite cells with small nucleus. The small infected cells are on the periphery of the foci but usually not immediately adjacent to the infected ECs. These observations provide several insights into the cell-to-cell virus progression. Because the immunofluorescence staining requires a certain level of viral protein accumulation in the cells, the technique is expected to preferentially detect late stages of infection. Therefore, by the time the infection is detectable in the primary infected cells, the secondary infections already occur in the neighboring cells. We speculate that the small satellite infected cells might have a faster replication cycle than the cells with large nuclei thus the immunofluorescence signal was detected earlier. Alternatively, the brighter immunofluorescence signals in the small cells might result from the more concentrated viral protein due to the smaller size of their cytoplasm.

**Figure 6.**
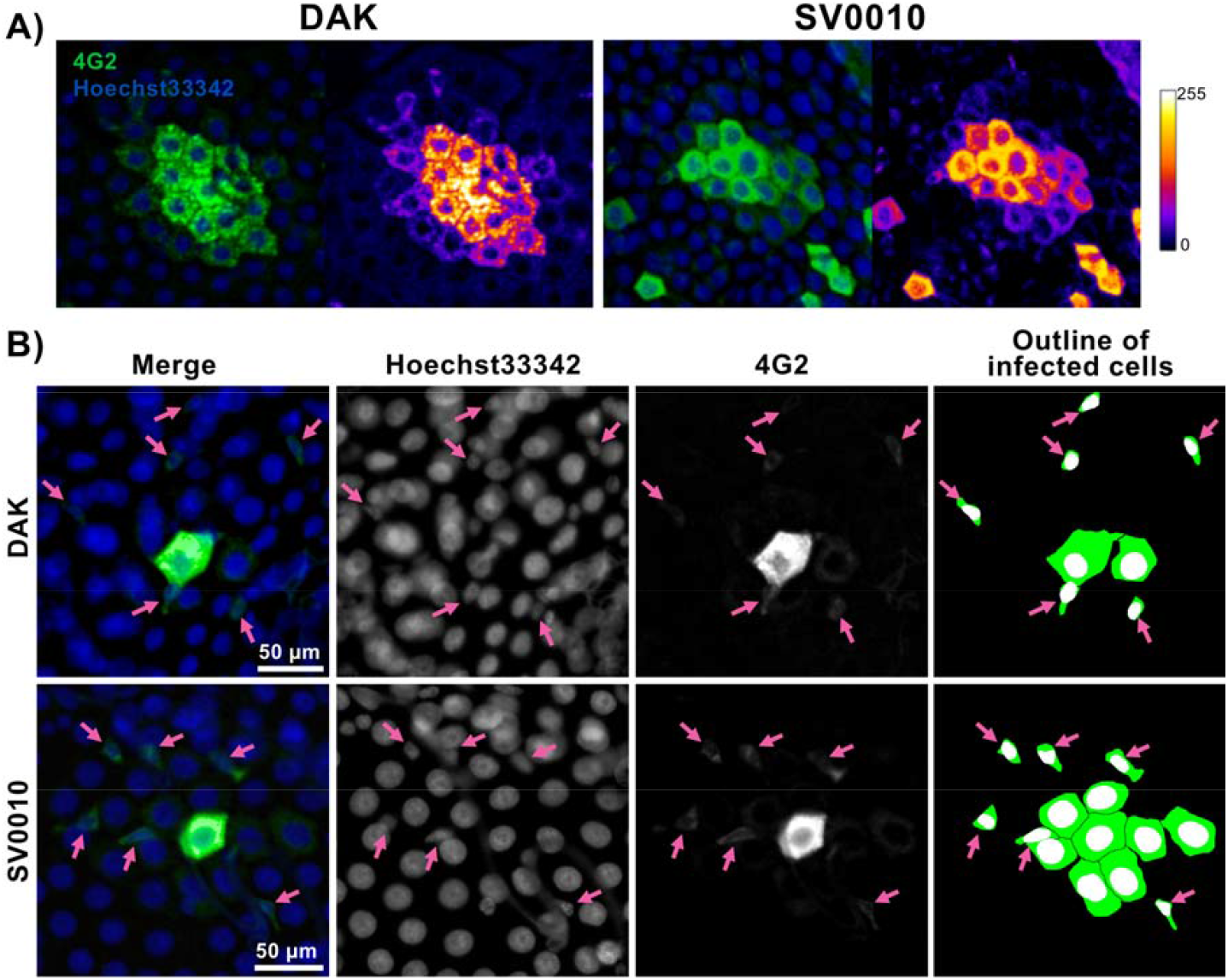
Progression of ZIKV infection in midgut cells. (A) Immunofluorescence staining of infected cell clusters with secondary infections. Immunofluorescence images of the African ZIKV strain (DAK) and the Asian ZIKV strain (SV0010). The heatmap demonstrates the green channel fluorescence intensity on a scale ranging from 1 to 255 arbitrary units. (B) infection foci of both ZIKV strains at 2 dpibm. Pink arrow heads indicate infection in small cells of the focus periphery.

## Discussion

Although it is well-established that the African ZIKV lineage exhibits higher transmissibility in *Aedes* mosquitoes compared to the Asian ZIKV lineage (Aubry et al., 2021; Calvez et al., 2018; Hery et al., 2019; Ou et al., 2021), limited knowledge exists regarding ZIKV infection kinetics during the early steps of mosquito infection and the virus’s interaction with various mosquito tissues. To enhance our understanding of the differential transmissibility the two ZIKV lineages, our study compared the intra-host tissue-level infection kinetics of African and Asian ZIKV strains in *Ae. aegypti* mosquitoes by examining both the infection intensity (amount of infectious virus as measured by plaque assay) and prevalence during midgut infection, dissemination, salivary gland infection, and virus release in saliva over time. We observed that the African ZIKV strain established a midgut infection, disseminated systemically, and was released in saliva more rapidly than the Asian ZIKV strain.

We then utilized *in situ* immunofluorescence staining to investigate cell-to-cell infection kinetics and pinpoint differential infection barriers between the two ZIKV strains. Although previous studies have employed similar immunofluorescence staining techniques to visualize the progression of arbovirus infection in mosquito tissues (Cui et al., 2019; Salazar et al., 2007), these studies lacked quantitative assessment of infection, hindering a comprehensive comparison of infection kinetics. By quantitatively analyzing cell-to-cell infection kinetics, we were able to provide insights into host-virus interactions at a higher resolution, contributing valuable information on the differential infection kinetics between the two ZIKV lineages. The key feature of African ZIKV is a more robust virus propagation which may be attributed to increased virus production in individual cells, faster viral spread within tissues, or both. In the midgut tissue, the African ZIKV strain established primary infections in the midgut epithelial cells, replicated in the primary infected cells, and spread throughout the mosquito tissue more efficiently than the Asian ZIKV strain.

Combining infectious virus detection by plaque assay and *in situ* immunofluorescence staining of the midgut, our data suggest that African ZIKV replicates at a faster rate than Asian ZIKV. At 1 dpibm, infectious Asian ZIKV could not be detected in any mosquito by plaque assay despite positive immunofluorescence signals in the midgut, suggesting either incomplete viral assembly within the first day or infectious titers below the limit of detection by plaque assay. Conversely, at the same time point, half of the African ZIKV-fed mosquitoes exhibited detectable infectious viruses, indicating a more robust propagation. The eclipse phase of Asian ZIKV replication in the midgut aligns with previous finding with the I-44 Mexican ZIKV strain (Cui et al., 2019). The faster replication of African ZIKV was also observed in mammalian cells such as human neural stem cells, primary human astrocytes, Vero (African green monkey kidney), HEK-293 (human embryonic kidney), and RK-13 (Rabbit kidney) cell lines (Hamel et al., 2017; Simonin et al., 2016; Smith et al., 2018) as well as several insect cell lines (Smith et al., 2018) suggesting that the phenotype is conserved across mammalian and insect hosts (reviewed in (Simonin et al., 2017)).

Using immunofluorescence staining of midguts at an early timepoint, we could estimate the number of primary infection foci. This analysis demonstrated a major bottleneck during the establishment of midgut infection shared by African and Asian ZIKV. With bloodmeal titers of 7 log_10_ PFU/mL, each mosquito is expected to ingest 10,000-20,000 PFU (1-2 μL of bloodmeal), yet the majority of midguts had a number of primary infected cells in the range of hundreds, meaning that only a few percent of infectious virions successfully establish a midgut infection. The scale of the bottleneck and the size of founding population are similar to previous estimates based on sequencing approaches for dengue virus (Lequime et al., 2016; Sim et al., 2015), and immunofluorescence staining-based techniques for dengue virus (Le Coupanec et al., 2017; Salazar et al., 2007) and Venezuelan equine encephalitis infectious clones (Forrester et al., 2012).

Because *in situ* immunofluorescence staining preserves spatial information of virus spread in mosquito tissues, the technique can be used to investigate characteristics of virus infection in different cell types, in addition to the rate of cell-to-cell spread. We observed that midgut cells with small nuclei support faster ZIKV replication than midgut cells with large nuclei. This observation suggests that different cell populations have a different ability to support virus replication. The identity of these small midgut cells has yet to be determined due to a lack of tools for identifying cell types, such as antibodies or transgenic reporter lines. A recent single-cell transcriptomics study that identified markers for each midgut cell population will make these molecular tools feasible in the future (Cui & Franz, 2020).

Another intriguing finding from our time-course analysis is that ZIKV infection in midguts exposed to a higher bloodmeal titer (7 log_10_ PFU/mL) progressed more quickly than in midguts exposed to a lower bloodmeal titer (5 log_10_ PFU/mL). Although it may seem logical that a larger virus inoculum would result in a more productive infection, our results demonstrated that a higher bloodmeal titer did not only result in a greater number of virus particles initiating the infection, but also a faster replication and spread in the mosquito tissues. This implies that the midgut responses differ according to the inoculum size. This is possibly because the higher amount of virus during the early infection could manipulate host machineries and immune system to favor virus replication. This is consistent with a previous study on dengue virus (Novelo et al., 2019), which observed that a higher bloodmeal titer resulted in lower viral loads in mosquito tissues even during the crossing of subsequent infection barriers at later timepoints.

Our findings of differential intra-host infection kinetics contribute to a better understanding of the differences in transmission efficiency between African and Asian ZIKV strains. Further investigation into the viral and host factors driving the differences in propagation efficiency between these two ZIKV lineages is essential for unraveling the underlying mechanisms and informing the development of ZIKV transmission control strategies. Advanced techniques such as CRISPR/Cas9 genome editing, proteomics, and single-cell RNA sequencing can be employed in future studies to provide more insights into the molecular determinants of the observed strain-specific infection kinetics.

## Acknowledgements

This work was supported by Thailand Program Management Unit for Human Resources & Institutional Development, Research and Innovation (PMU-B), NXPO, grant number B17F640002 to NJ, and the French Government’s Investissement d’Avenir program Laboratoire d’Excellence Integrative Biology of Emerging Infectious Diseases (grant number ANR-10-LABX-62-IBEID) to LL. The personnel exchange for the collaboration was supported by the European Union’s Horizon 2020 research and innovation programme under the Marie Skłodowska-Curie grant agreement No 734486 (SAFE-Aqua). The 4G2 antibody was a gift from Dr. Bunpote Siridechadilok (BIOTEC). The African ZIKV DAK AR 41524, NR-50338 was obtained through BEI Resources, NIAID, NIH, as part of the WRCEVA program. The Asian ZIKV SV0010/15 was obtained from the Armed Forces Research Institute of Medical Sciences (AFRIMS) and the Department of Disease Control, Ministry of Public Health through the Cluster Program Management Office, NSTDA.

## Supplementary Figures

**Supplementary Figure S1.**
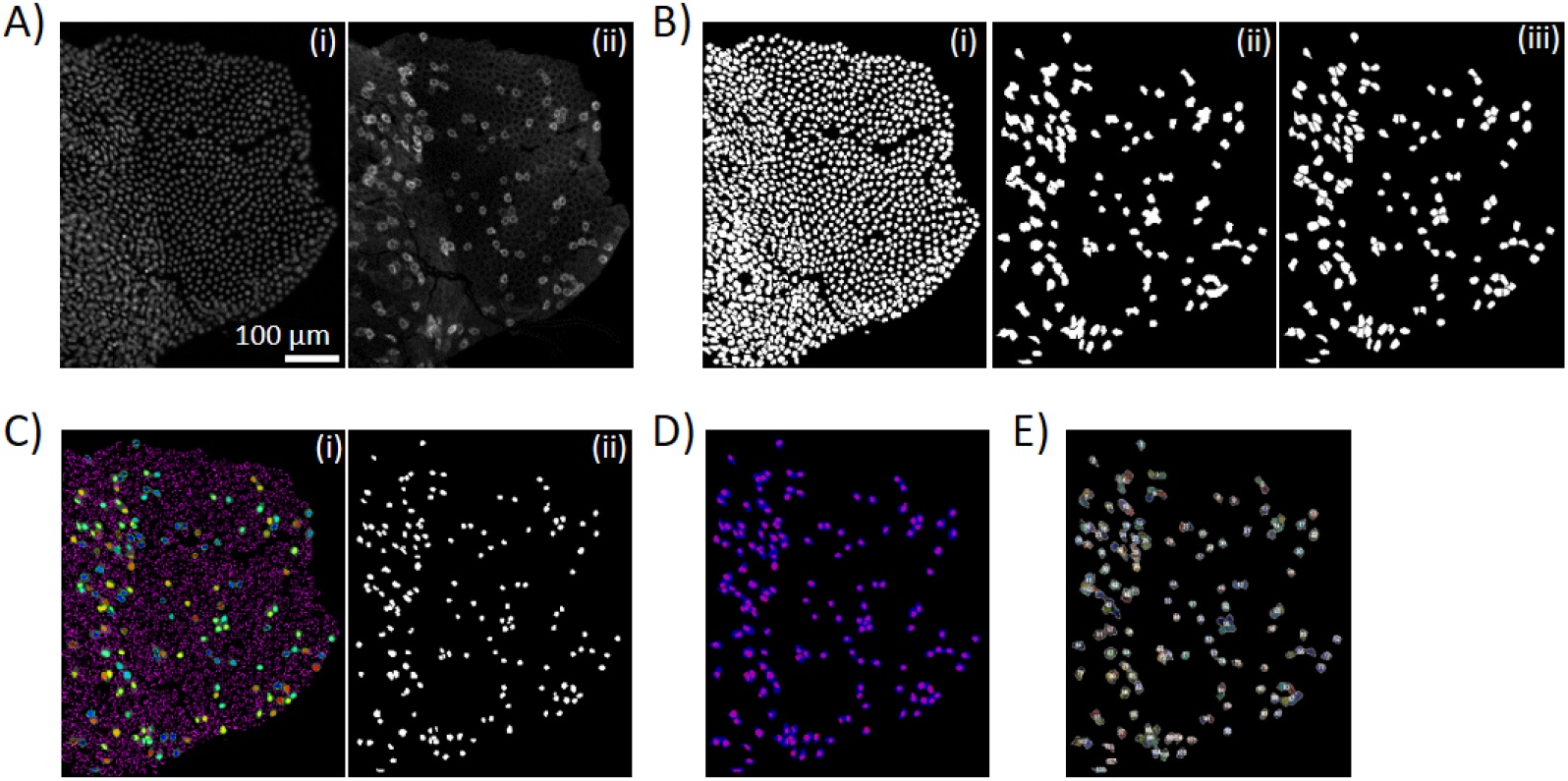
Detection of infected foci/cells and their nuclei. A) Raw image from Hoechst33342 (i) and Alexa 488 (ii, ZIKV E proteins) channels. (B) Nuclei (i) of midgut cells and infection foci (ii) identified by Stardist nuclear segmentation and trainable Weka segmentation, respectively. Individual infected cells (iii) were further separated using watershed and manual segmentation. (C) Infection foci mask (i) was used for extracting nuclei of the infected cells (ii). (D) An overlay image of infected foci and their nuclei. (E) Final processed image showing infection foci boundaries (light gray line) and their nuclei (color).

**Supplementary Figure S2.**
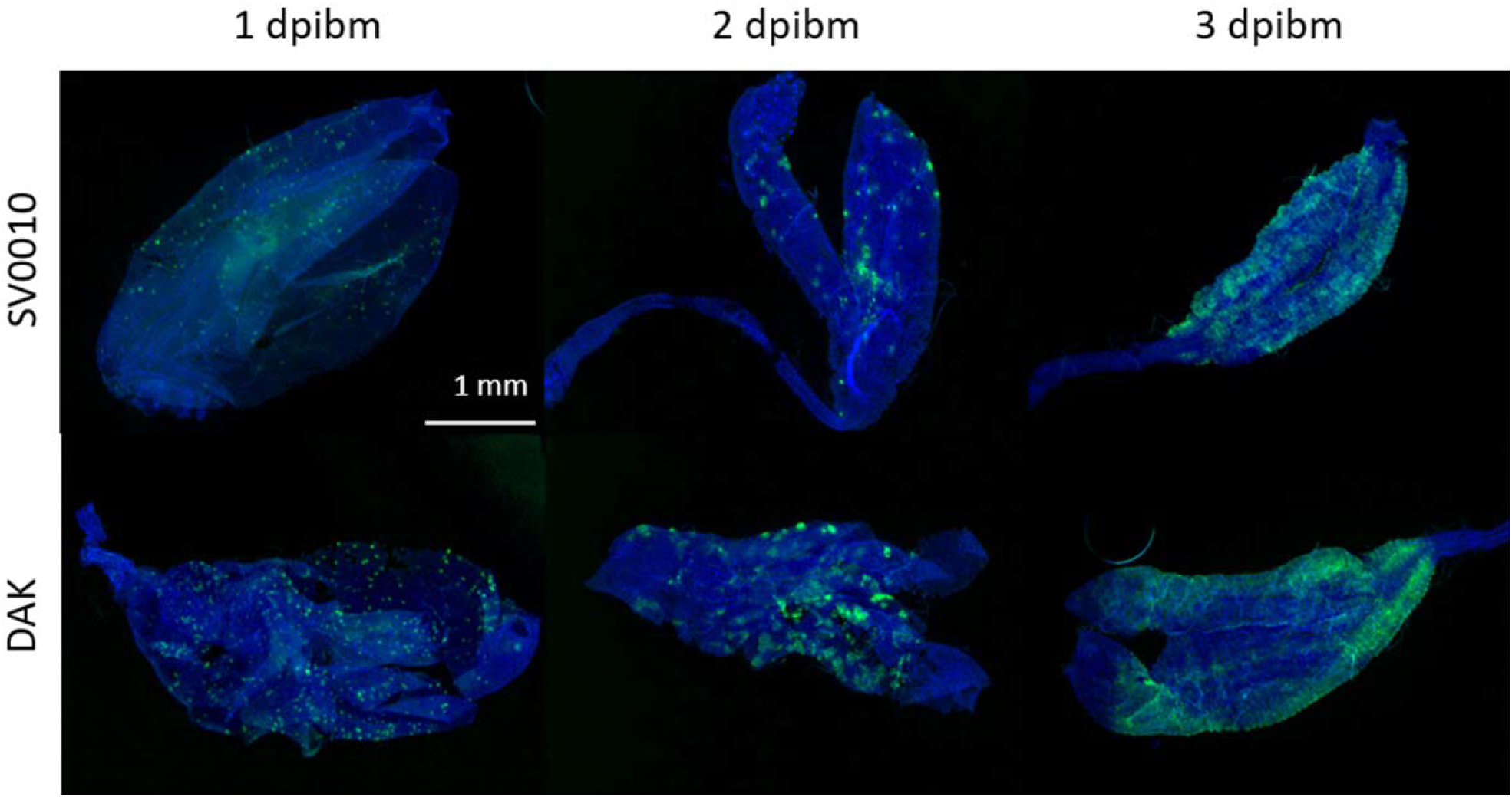
Representative images of infection kinetics at 1, 2, and 3 dpibm of midguts infected by 7 log_10_ PFU/mL of the African ZIKV strain (DAK) and the Asian ZIKV strain (SV0010).

